# Functional reorganization of brain regions supporting non-adjacent dependency learning across the first half year of life

**DOI:** 10.1101/2024.04.03.587880

**Authors:** Lin Cai, Yoko Hakuno, Masahiro Hata, Ei-ichi Hoshino, Takeshi Arimitsu, Naomi Shinohara, Takao Takahashi, Stuart Watson, Simon Townsend, Jutta L. Mueller, Yasuyo Minagawa

## Abstract

Pre-babbling infants can track nonadjacent dependencies (NADs) in the auditory domain. While this forms a crucial prerequisite for language acquisition, the neurodevelopmental origins of this ability remain unknown. We applied functional near-infrared spectroscopy in neonates and 6-7-month-old infants to investigate the neural substrate supporting NAD learning using tone sequences in an artificial grammar learning paradigm. Detection of NADs was indicated by left prefrontal activation in neonates while by left supramarginal gyrus (SMG), superior temporal gyrus (STG), and inferior frontal gyrus activation in 6-7-month-olds. Functional connectivity analyses further indicated that the neonate activation pattern during the test phase benefited from a brain network consisting of prefrontal regions, left SMG and STG during the rest and learning phases. These findings suggest a left-hemispheric learning-related functional brain network may emerge at birth and be strengthened by complex auditory input across the first half year of life, providing a neural basis for language acquisition.

## Introduction

Humans are born with sophisticated auditory abilities, possibly shaped by prenatal experience and a relatively mature auditory system [1, 2]. Studies with newborn infants have thus demonstrated impressive abilities in the auditory domain including discrimination of stimuli based on various auditory features [3, 4], auditory learning of novel sounds [5, 6] and computation of more complex sound sequences [7–10]. The latter is of particular relevance for language acquisition, as human syntax relies on the ability to decode a hierarchical structure from sequentially organized auditory input. The sequential auditory input can involve more or less complex dependency patterns ranging from simple adjacent dependencies, i.e., the relation of two consecutive stimuli, to multiple embedded non-adjacent dependencies. Non-adjacent dependencies (NADs) are important for language because they allow the meaningful relation of elements across a distance which allows the formation of complex and hierarchical sentence structures [11]. For example, the sentence “The baby smiles”, which contains a NAD between the verb suffix –s and the noun, can be extended to “The baby who is sitting on her mother’s lap smiles” which contains a further NAD between the suffix –ing and the auxiliary “is”, creating a nested structure of NADs. In order to analyze such sentences, the dependent elements have to be stored and retrieved across variable distances. The basic computational mechanisms underlying the ability to detect NADs have been studied both using linguistic as well as non-linguistic materials [12–15]. The ability to learn NADs seems to be present from early on, both when encoded in speech [16] and when encoded in computationally simpler sine-tone sequences [12, 13]. Even non-human animals seem to be able to detect NADs in some cases [17, 18] suggesting that the basic ability may not be unique to humans and constitutes an important computational basis for language which is present early in development and potentially rooted in our primate ancestors. What is not known, though, is i) from which age onwards the ability to detect NADs is present in humans and ii) which brain areas are involved in this ability in those early developmental stages.

The ability to learn NADs in the auditory domain and its developmental trajectory have been subject to intense investigations in infants and young children [11, 19].

Previous behavioral investigations have shown that a sensitivity to NADs, embedded in natural or artificial language, appears after the first year of life [20–23] (around 15-19 months), and the ability to discriminate nonadjacent repetitions (e.g., ABA) from other patterns (e.g., ABB or ABC) is present in 5- to 7-month-old infants [24, 25]. However, such findings may be constrained by the restrictions of behavioral measures, i.e., the dependence of those measures on overtly observable responses. Electrophysiological methods, for example, present a more fine-grained approach toward directly probing the recognition of NADs, and have correspondingly validated infants’ earlier sensitivity to NADs [15, 26]. For instance, an event-related potential (ERP) study using the familiarization-test paradigm suggests that 4-month-old monolingual infants can passively learn NADs from a foreign language, indicated by a late positive ERP effect [15]. Several other electrophysiological studies attested NAD learning before the first birthday [12, 26, 27]. While the learning of adjacent dependencies has been demonstrated already in newborns [9, 10], there is no evidence so far for the learning of NADs in newborns.

Unfortunately, electrophysiological evidence alone cannot provide precise information concerning the question of which brain regions are involved in NAD learning. Recent functional magnetic resonance imaging (fMRI) advances in adults reveal that Broca’s region is involved in the processing of NADs [28–30]. Similarly, van der Kant et al., (2020) [13] found, using functional near-infrared spectroscopy (fNIRS), that NAD violation detection was subserved by a left-hemispheric temporo-fronto-parietal network for linguistic stimuli in 2-year-olds. On the other hand, 3-year-olds, but not 2-year-olds were able to learn NADs consisting of non-linguistic tone stimuli engaging a bilateral temporo-parietal network. However, tone stimuli do not seem to be associated with later NAD learning per se as demonstrated by a recent ERP evidence showing that infants who were only 5 months of age can track embedded NAD structures between simple sine tone stimuli [12]. A study focusing on the specific role of prosodic cues for NAD learning in 9-month-old infants revealed the contribution of frontotemporal brain regions in the more difficult, monotonous condition and the engagement of temporal brain areas in the presence of prosodic cues [14]. Yet, this study leaves unclear whether the infants learned NADs between two specific elements or only positional regularities. Together, these findings indicate that NAD learning may be present early and that neural networks that include classic language areas are involved in those learning processes from early childhood onwards, at least in the case of linguistic stimuli. However, it is not known whether similar neural networks support NAD learning and processing from birth and how they develop across the first months of life.

A growing body of research demonstrates that neonates possess an adult-like ventral pathway for language, which connects the anterior temporal lobe with the ventrolateral prefrontal cortex by the extreme capsule [31]. Likewise, the part of the dorsal pathway connecting the temporal cortex to the premotor cortex is also already present at birth.

These structural connections may serve as a neural basis for rule learning [31], maternal speech perception [32], and phonological learning [5] in neonates. By contrast, the part of the dorsal pathway connecting the temporal cortex to Broca’s area develops much later, but it catches up during the first post-natal months [33]. Note that this connection has been argued to be involved in parsing more complex syntactical structures during childhood [31, 34]. Taking into account both the early functionality of the prefrontal cortex and its relative immaturity, a very recent review [35] proposed that infants learn actively from their environment early on and that this is supported by their (anatomically) immature prefrontal cortex. Taken together, these studies reveal that an intriguing functional brain network consisting of several distributed key brain regions for language processing has emerged since birth. Further, they suggest that the remarkable learning ability of NADs in older infants reported above could be supported by the ventral and dorsal pathways despite the relative immaturity of the prefrontal cortex (and its dorsal connection to posterior brain areas).

Because NAD learning seems to be present early in development and distributed across various vertebrate species and also relevant for core processes of language, as outlined above, we hypothesize that i) humans are born with the ability to track NADs, and ii) the brain regions that support this process may include areas that are also involved in early language processing including left hemispheric fronto-temporal networks. In the current study, we examined the neurodevelopmental origins of NAD learning using non-linguistic tone sequences in order to further understand the ontogenesis of human language. Specifically, we adopted the fNIRS technique to test neonates (1-5 days old) in Experiment 1 and infants (6-7 months old) in Experiment 2 using an auditory artificial grammar learning paradigm (Fig 1B). In both experiments, the stimuli were composed of a series of perceptually simple tone sequences, each of which comprised three frequency-modulated sine tones from six categories of pitch contour (A, B, C, D, X1, X2; see Fig 1A). Each sine tone category had various pitch variants which were used to avoid stimulus-specific learning. In the learning (Learning) phase, participants were exposed to 60 standard tone triplets conforming to NAD rules (e.g., AXB or CXD grammar) for approximately 6 minutes. Subsequently, during the test (Test) phase, they were presented with the familiar standard sequences and sequences comprised of novel pitch variants of the acoustic categories heard during the Learning phase arranged into either ‘Correct’ triplets (e.g., AXB or CXD) or ‘Incorrect’ triplets (e.g., AXD or CXB), thereby testing whether individuals were able to generalize the NAD rules from the Learning phase and detect violations. In Experiment 1, we additionally recorded the hemodynamic activities of neonates during the pre-task resting-state (Pre-Rest) phase, the Leaning phase and the post-task resting-state (Post-Rest) phase. Pre-Rest and Post-Rest phases were inserted before and after Learning and Test phases, respectively. The two resting-state phases allowed us to examine changes in functional connectivity (FC) from Pre-Rest to Learning phases or from Pre-Rest to Post-Rest phases, as well as the relationship between the FC changes and the activation to NAD violations during the Test phase (Fig 1C). In Experiment 2, we exposed 6-7-month-olds to Learning and Test phases without resting-state phases but we performed fNIRS scans only during the Test phase (Fig 1C). The decision to limit scan time in this experiment was based on the increased level of physical activity and thus potentially decreased compliance with the procedure in this age group.

**Fig 1.**
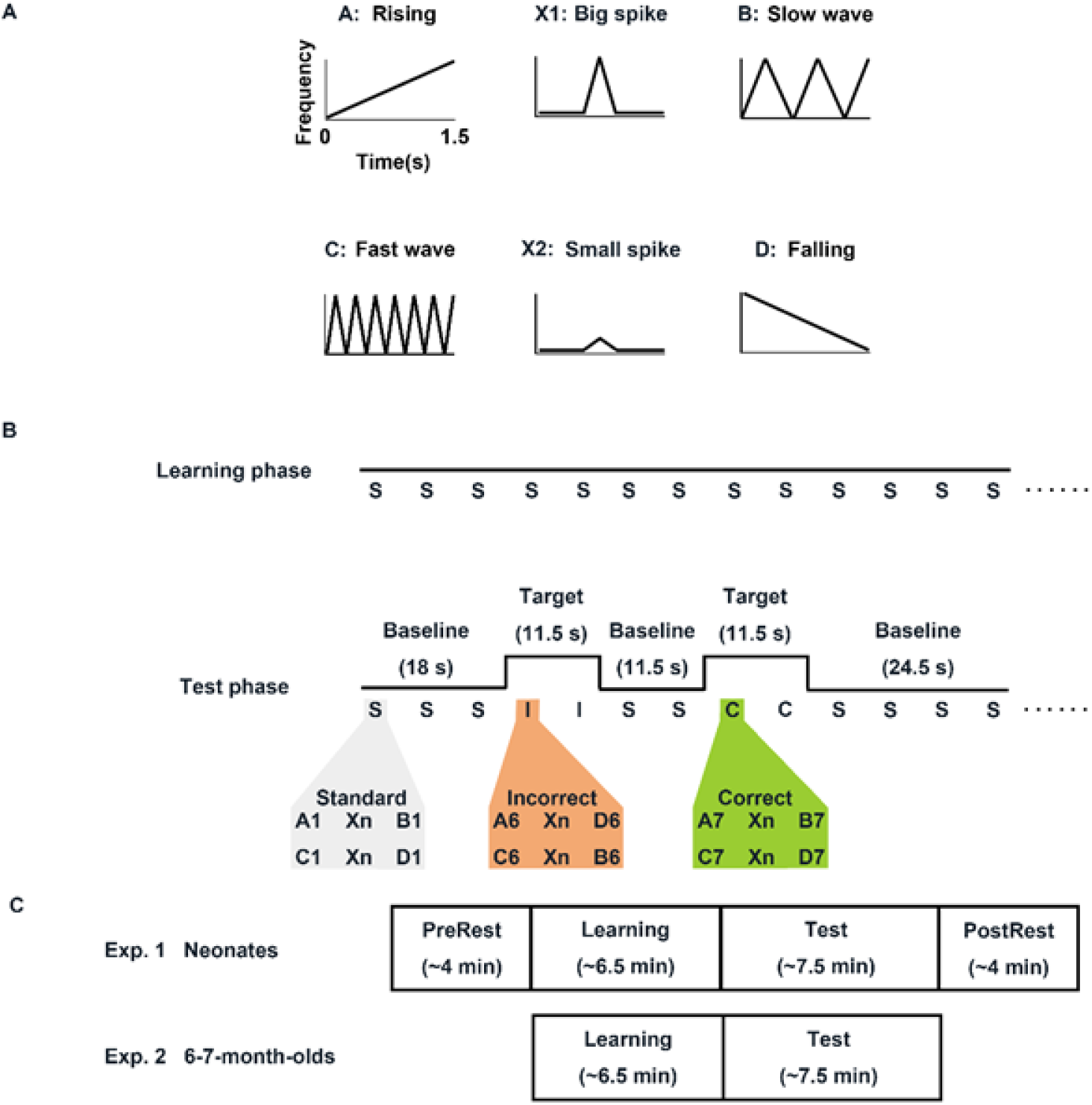
Stimuli and experimental design. (A) The pitch contours of six acoustic categories. **(B)** Experimental design. In the Learning phase, standard triplets were repeated 60 times. In the Test phase, 2 types of target trials (Correct and Incorrect conditions) were separated by a jittering baseline condition in which the same standard triplets with the Learning phase were presented. **(C)** Measurement phases for two experiments.

Thus, the first aim of Experiment 1 was to reveal whether human neonates can extract NAD relations and generalize them to novel tone sequences, which would be indicated by a greater neural response to incorrect sequences than to correct ones during the Test phase. The second aim of Experiment 1 was to identify, in the case of successful NAD violation detection, the brain networks underlying NAD learning at birth by examining FC changes (Learning minus Pre-Rest or Post-Rest minus Pre-Rest) and their correlations to cerebral responses to learned NAD relations as measured during the Test phase. The aim of Experiment 2 was to examine what brain networks underlies NAD learning in 6-7-month-old infants. By combining these two experiments, we could delineate a picture of emergence and development of functional brain network underlying NAD learning across the first half year of life.

## Results

### Experiment 1

As illustrated in Fig 2A (left and middle), mean hemodynamic response within the 5-s time window to the Correct and Incorrect conditions as indicated by oxygenated hemoglobin (HbO) changes was compared with that of the baseline period for each channel. Permutation t-tests (SI Appendix, Table S4) confirmed that neonates showed significantly decreased activation for the Correct condition compared to baseline.

**Fig 2.**
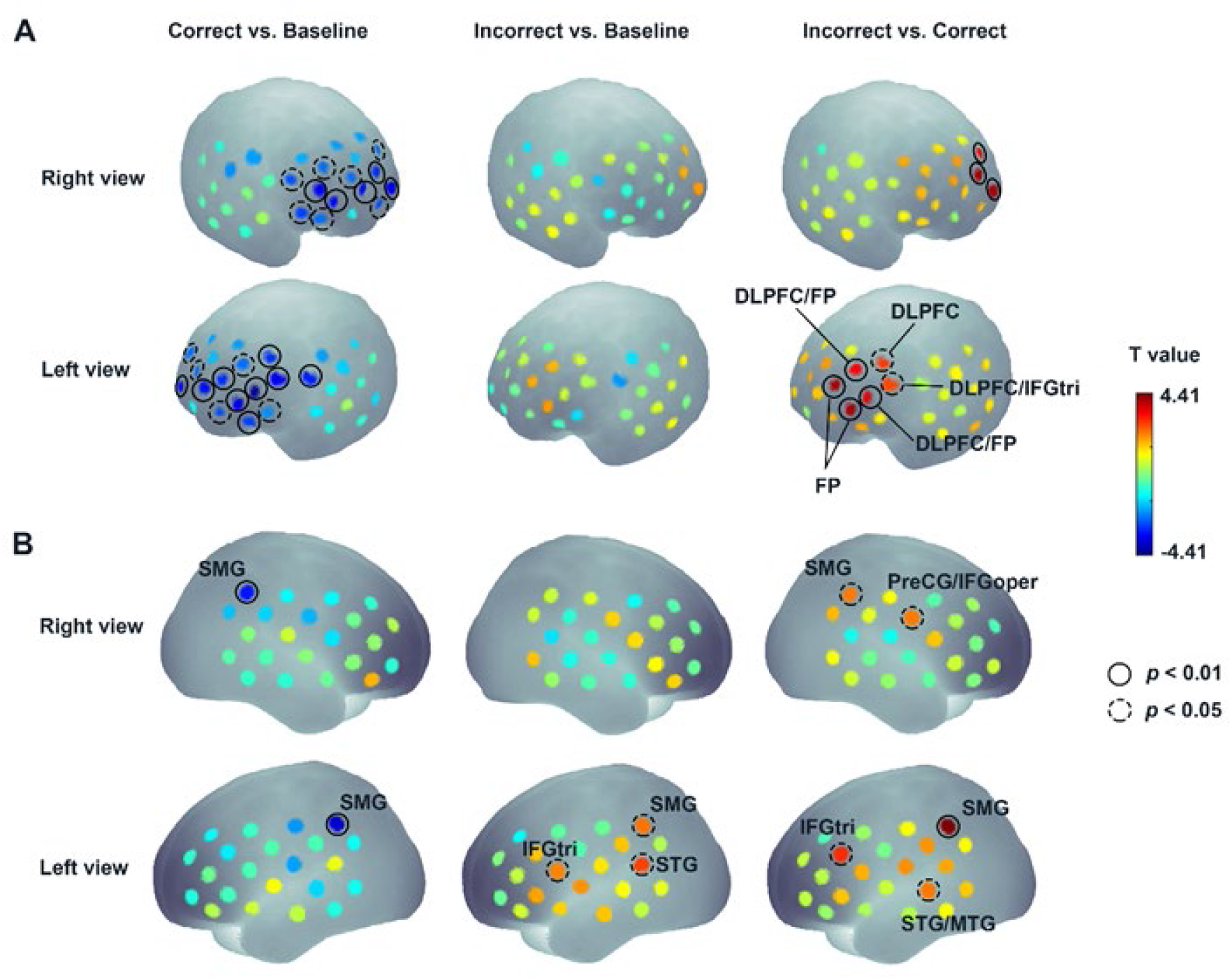
The statistically significant effects in the channel-by-channel permutation t- test analysis. (A) Experiment 1: neonates. **(B)** Experiment 2: 6-7-month-olds. Left, middle and right panels show brain regions that were activated significantly for three condition contrasts of Correct vs. Baseline, Incorrect vs. Baseline, and Incorrect vs. Correct, respectively. The circles denote significant channels (solid circles: permutation p < 0.01; dotted circles: permutation p < 0.05).

Significant activation was chiefly found in the prefrontal regions including channel (Ch) 3, Ch 25-26, Ch 28-39, and Ch 41-43 (Fig 2A, left). On the other hand, although no significant differences were observed for the Incorrect vs. baseline (Fig 2A, middle), a comparison between the Incorrect and Correct conditions (Fig 2A, right) revealed a greater activation dominantly in the left hemisphere. These significant brain regions included left (L) frontal pole (L-FP) (Ch 30: t = 4.05, p = 0.001; Ch 35: t = 4.04, p = 0.001), L dorsolateral prefrontal cortex (L-DLPFC)/L-FP(Ch 34: t = 3.49, p = 0.003; Ch 39: t = 3.32, p = 0.003), L-DLPFC/ L triangular part of inferior frontal gyrus (L-IFGtri) (Ch 38: t = 2.71, p = 0.012), and L-DLPFC (Ch 43: t = 2.84, p = 0.008). (See SI Appendix, Table S4 for a summary of statistical results and SI Appendix, Fig S2 for the grand averaged time courses of the hemodynamic responses for two experimental conditions)

Experiment 1 further explored the FC patterns during the Pre-Rest phase, the Learning phase, and the Post-Rest phase. For each of the three phases alone, there were a large number of significantly stronger FCs relative to a respective zero baseline, with more than 445 significant FCs (Bonferroni corrected p < 0.05, SI Appendix, Fig S4). To further explore the locations of the most significant FCs in the brain, we set stringent thresholds at four levels (Bonferroni corrected p < 5e-6, p < 5e-7, p < 5e-8, and p < 5e-9) as indicated in Fig 3A. It showed that the Learning phase exhibits a larger number of significant FCs compared to the other two resting-state phases. Particularly, FCs connecting frontal and temporal regions emerged during the learning phase, which were absent during the other two resting-state phases. Furthermore, we compared FC differences between the Learning and Pre-Rest phases, as well as between the Post-Rest and Pre-Rest phases. For the first comparison (Learning minus Pre-Rest) we found 96 significantly increased FCs (uncorrected p < 0.05; Fig 3B, Left). For the second comparison we found 13 significantly increased FCs and 6 significantly decreased FCs (uncorrected p < 0.05; Fig 3B, Right).

**Fig 3.**
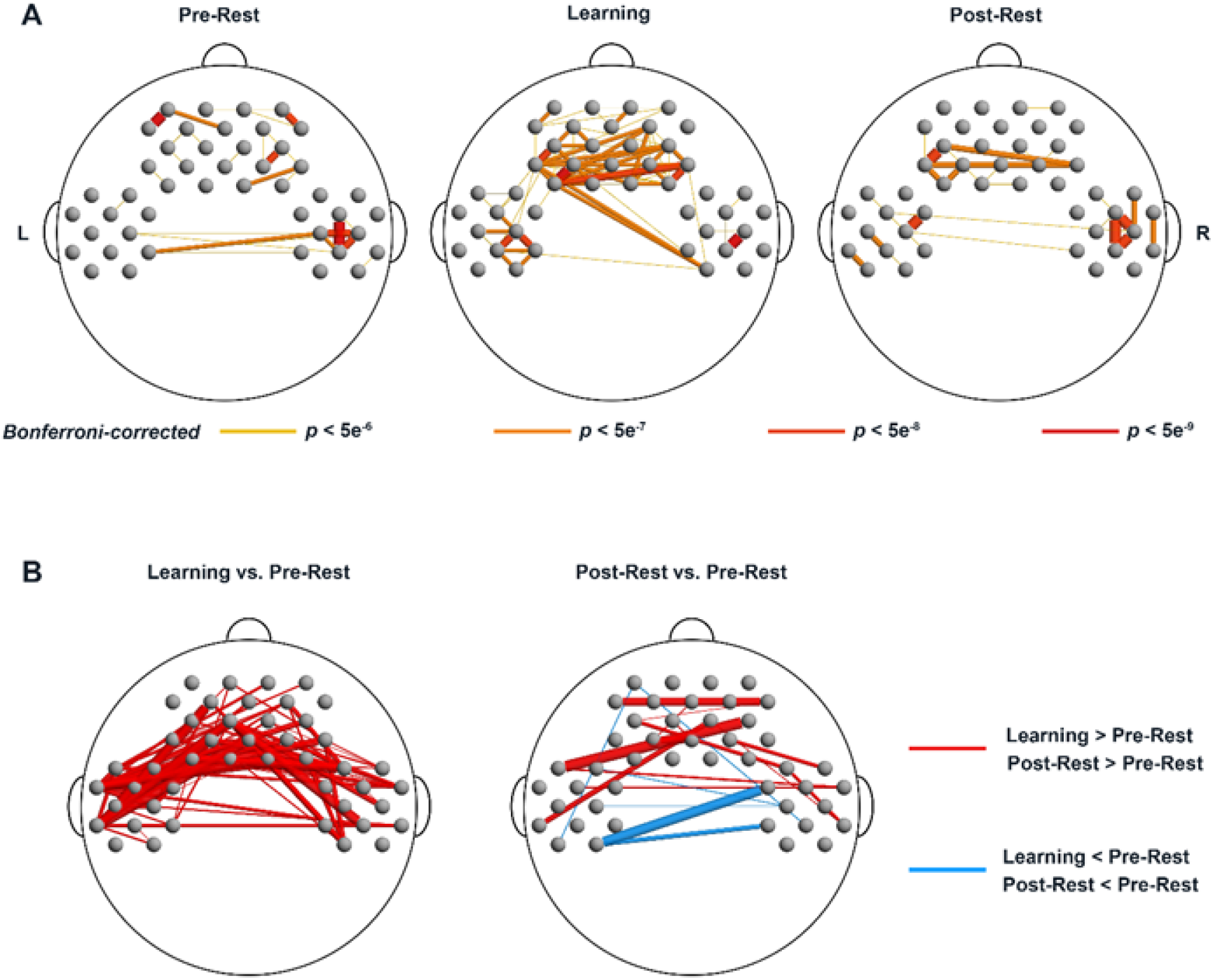
The FC patterns in Experiment 1: neonates. (A) The most significantly increased FCs for three phases against a respective zero baseline. Different colors and line thickness indicate different thresholds. Both darker color and thicker line indicate the more stringent threshold. **(B)** The FCs with significant changes for the contrasts of Learning phase vs. Pre-Rest phase, and Post-Rest phase vs. Pre-Rest phase.

To examine how FCs during NAD learning relate to actual learning success, we analyzed the correlations between the strength of FCs with significant changes during the Learning phase (i.e., Learning minus Pre-Rest) and the degree of activation in those 6 prefrontal channels that were significantly activated during the Test phase (Ch 30, Ch 34, Ch 35, Ch 38, Ch 39, Ch 43, termed seed channels from here onwards). Thus, FCs connecting any one of those 6 seed channels were examined. Notably, as illustrated in Fig 4A, the strength of 9 FCs during the Learning phase (Ch 30-Ch 9, r = -0.52; Ch 34- Ch 9, r = -0.66; Ch 35-Ch 9, r = -0.55; Ch 38-Ch 17, r = -0.54; Ch 39-Ch 40, r = -0.68; Ch 43-Ch 2, r = -0.57; Ch 43-Ch 17, r = -0.56; Ch 43-Ch 19, r = -0.60; Ch 43-Ch 22, r = -0.56; all ps < 0.05) were negatively correlated with the activations in seed channels (A canonical negative relationship sees Fig 4B left). From the perspective of anatomy, these FCs mainly connected prefrontal regions to L supramarginal gyrus (L-SMG) and L superior temporal gyrus (L-STG), as well as right (R)-IFG, R precentral gyrus (R-PreCG), and R-STG. To further examine whether these 9 FCs during the Learning phase (Fig 4A) and other FCs within these channels form a learning-related brain network together, we analyzed the correlations between the FC strength of any two non-seed channels involved in the 9 FCs (i.e., L-STG, L-SMG, R-IFG, R-STG, R-PreCG, DLPFC) and the activation degree of 6 prefrontal seed channels. As shown in Fig 4B (right), the strength of FC linking L-SMG (Ch 2) and L-STG (Ch 9) was significantly negatively related to the activation degree of all 6 seed channels (r values: Ch 30, -0.68; Ch 34, -0.81; Ch 35, -0.70; Ch 38, -0.58; Ch 39, 0.67; Ch 43, -0.66, ps < 0.05). Finally, all 10 (9+1) intrinsic FCs showing negative correlations with activation in 6 prefrontal channels formed a learning-related brain network as shown in Fig 4C, which may be responsible for NAD learning.

**Fig 4.**
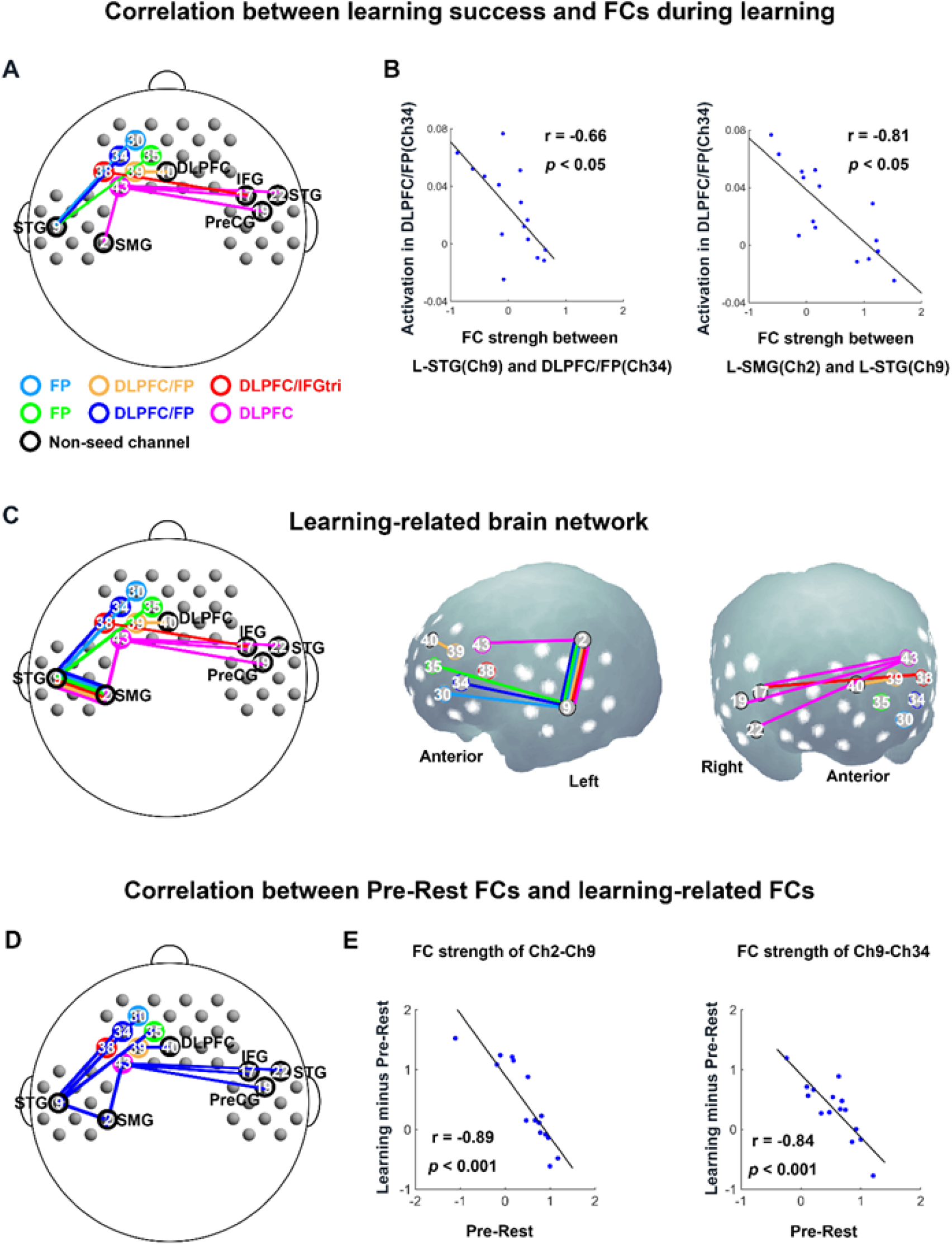
Functional brain network underlying NAD learning in Experiment 1: neonates. (A) Relations between prefrontal activation during the test phase (learning success) and FCs with significant changes from the Pre-Rest phase to the Learning phase (FCs during learning). Seed channels are indicated by circles with 6 different colors and non-seed channels indicated by black circles. The lines with 6 different colors represent FCs displaying significantly negative correlation with activations in seed channels. **(B)** The negative correlation between activation in Ch 34 and FC strength (Ch 9 - Ch 34; Ch 2 - Ch 9). The blue dots represent data from 15 neonates. **(C)** 2D and 3D maps of the learning-related brain network, where the strength of all FCs was negatively correlated with activation in the 6 seed channels. **(D)** Relations between FC strength during the Pre-Rest phase and FC strength changes from the Pre-Rest phase to the Learning phase (Learning minus Pre-Rest). Blue lines indicate negative correlations. **(E)** The scatter plots from two canonical negative correlations in Fig 4D. The blue dots represent data from 15 neonates.

To better understand the reason why the learning-related brain network with 10 intrinsic FCs was negatively related to the prefrontal activation found for NAD violation detection, we explored the relationship between FC strength during the Pre-Rest and Learning phases for the 10 intrinsic FCs. We found 9 (out of 10) negative correlations between FC strength during Pre-Rest and Learning minus Pre-Rest to be significant (r values: Ch2-Ch9, -0.89; Ch9-Ch30, -0.71; Ch9-Ch34, -0.84; Ch9-Ch35, -0.80; Ch39-Ch40, -0.84; Ch2-Ch43, -0.82; Ch17-Ch43, -0.58; Ch19-Ch43, -0.68; Ch22-Ch43, -0.76, ps < 0.05) (Fig 4D and 4E). This meant that while most neonates showed stronger FCs during the Learning phase compared to the Pre-rest phase (SI Appendix Fig S5), neonates with stronger FC as a default state (i.e., Pre-Rest) tended to show less increment of FC from Pre-Rest to Learning phases. Likewise, a negative correlation between FC strength for Pre-Rest and that for Post-Rest minus Pre-Rest (Ch34-Ch17) was confirmed, which revealed a similar link between default-state and later observed changes (See SI Appendix, Fig S6).

### Experiment 2

As illustrated in Fig 2B (left and middle), the mean hemodynamic response within the 5- s time window to the Correct and Incorrect conditions as indicated by HbO changes was compared with that of the baseline period for each channel. Permutation t-tests (SI Appendix, Table S5) revealed significantly decreased responses to the Correct condition relative to the baseline in L-SMG and R-SMG (Ch 4: t = -3.71, p = 0.002; Ch 23: t = - 3.13, p = 0.003). For the Incorrect condition vs. baseline, significantly increased responses were exclusively observed in the left hemisphere including L-SMG (Ch 4: t = 2.32, p = 0.031), L-IFGtri (Ch 11: t = 2.17, p = 0.045), and L-STG (Ch 13: t = 2.75, p = 0.013). Notably, comparisons between the two conditions (Fig 2B, right) showed greater responses to the Incorrect condition relative to the Correct condition, particularly in the L-SMG (Ch 4: t = 4.41, p = 0.001), L-IFGtri (Ch 6: t = 2.95, p = 0.011), and L-STG/L middle temporal gyrus (L-MTG) (Ch 17: t = 2.28, p = 0.024). Other significant brain regions included R-SMG (Ch 23: t = 2.25, p = 0.046) and R-PreCG/R opercular part of inferior frontal gyrus (R-IFGoper) (Ch 29: t = 2.25, p = 0.040). (See SI Appendix, Fig S3 for the grand averaged time courses of the hemodynamic responses for two experimental conditions).

## Discussion

The current study investigated the neurodevelopmental origins of NAD learning by monitoring infants’ neural responses to NAD processing across two fNIRS experiments using an artificial grammar learning paradigm. In Experiment 1, we observed neonates showing significant activation differences between correct and incorrect NAD exemplars after a short Learning phase in left prefrontal regions mainly including DLPFC and FP. Despite no primary activation in either the anterior- or posterior-language areas, the degree of activation in prefrontal regions was negatively associated with the strength of FCs linking prefrontal regions and the posterior-language area during the Learning phase. Additional analyses revealed that the direction of this link may have been driven by the connectivity strength measured during the pre-task resting-state period. In Experiment 2, we found significant activation differences between the Correct and Incorrect NAD conditions in 6-7-month-old infants in mainly left-hemispheric brain areas including the left SMG, STG, and IFG with some activations in the right SMG. These findings demonstrate that both neonates and 6-7-month-olds can discriminate between correct and incorrect NADs in non-linguistic contexts after a brief Learning phase.

However, the underlying mechanisms are supported by partly different neural processes. Our results first suggest that even within the first half year of life there is a development toward the use of regions that are part of classical language networks for NAD processing. Second, the FC data demonstrate the involvement of classical language areas (SMG and STG) during NAD learning right after birth, and are suggestive of a gradual strengthening of the NAD-related brain network as infants are exposed to grammar-like input. Finally, such a brain network underlying the NAD processing for older infants seemed to come from the pre-established FC between prefrontal areas and posterior language areas at birth.

### The prefrontal cortex underlies neonates’ NAD learning

Experiment 1 with neonates shows activation in left prefrontal regions, including DLPFC and FP and IFG. Many prior studies have demonstrated that infants are active learners and, surprisingly, rely more on the prefrontal cortex to support their learning than previously thought [35]. For instance, two fNIRS studies demonstrated that sleeping 3- month-olds not only detect the violation of the learned rules [36] but also the occurrence of a novel sound [37] through employing the lateral prefrontal cortex. Even neonates’ left IFG seems to be involved in learning speech sounds [5] as well as regularities between syllables, specifically, repetition patterns [38]. Further, sleeping neonates recruited FP and IFG which were connected to the posterior-language areas in response to stimulation of maternal speech [32]. Another recent fMRI study also revealed that the prefrontal cortex may be sufficiently developed during young infancy to support stimulus-driven attention [39]. This evidence from the prefrontal cortex during early infancy is in line with our proposal that neonates’ brains are already sensitive to violations of NADs occurring in novel exemplars of pre-learned tone sequences.

Furthermore, the additional activation of left FP in neonates possibly reveals that FP serves as a functional ‘add-on’ at the apex of the hierarchy of lateral prefrontal processes, since FP activations are always concurrently observed together with lateral prefrontal activations [40]. Furthermore, the activation in a small part of IFG (Ch 38) might reveal that the neural underpinnings of processing NADs start involving the anterior-language area.

### Functional networks linked to learning effect in neonates’ prefrontal cortex

Interestingly, neonates did not show activation in the posterior-language areas (i.e., STG and SMG) as observed in 6-7-month-olds. However, negative relations between the degree of response in left DLPFC/FP and the strength of FCs linking DLPFC/FP and STG/SMG during the Learning phase indicate a potential neural mechanism including temporo-parietal cortices. Firstly, consistent with previous studies [41, 42], neonates showed a large number of FCs either during resting-state or stimulation phases, reflecting active sleep brain states of neonates [43]. However, among those phases, FCs during the Learning phase were stronger than during the Pre-Rest phase, both shown in long-range FCs between anterior-posterior brain regions and between hemispheres. In addition, consistent with findings from 3-month-old infants [41] and adults [44–46], the strength of FCs in the Post-Rest phase was higher or weaker than that in Pre-Rest phases. The FC difference between Pre-Rest and Post-Rest phases indicates that the exposure to sound stimuli during Learning and Test phases shaped resting-state brain networks of neonates, in particular, long-range FCs.

Most interestingly, when exploring the possible mechanisms of the negative learning-related brain network, we found that larger NAD learning effects in the prefrontal regions were linked to smaller increments of the left anterior-posterior FCs during the Learning phase compared to the Pre-Rest phase. We interpret this finding in the following way: Neonates equipped with stronger connections as a default state (Pre-Rest) for the learning-related network require less cognitive effort to strengthen the network in tracking NADs during the Learning phase, and such facilitated connections may have resulted in enhanced learning performance as reflected in the prefrontal activity measured during the Test phase. In contrast, for those neonates with weaker connections as a default state, a strengthening of the long-range anterior-posterior FCs is needed, as reflected by higher Learning minus Pre-Rest FCs during the Learning phase, but the still inefficient network may have led to weaker learning effects in the Test phase.

The learning-related brain network observed here involved the posterior-language areas including SMG and STG (see Fig 4). This suggests that neonates, despite showing exclusively prefrontal activation for the detection of NAD violation, relied on language-related brain regions during learning. The functional link between those regions may become more stable over time. The involvement of a larger left-hemispheric network for violation detection in 6–7-month-olds makes this seem plausible. Beyond language-related functions, the observed functional network could also serve more domain general purposes. Specifically, we speculate that the long-range FCs linking DLPFC and the inferior parietal cortex (e.g., SMG) reflect an early functionality of the already instantiated frontoparietal network, which is engaged in maintaining and manipulating information, since birth [47]. This fronto-parietal network could well play an important role in the service of NAD learning in neonates.

Although neonates tracked NADs by involving different brain areas compared to 6-7- month-olds, the findings on FC analyses revealed that neonates recruit functional brain networks that are probably supported by two structurally immature, yet to some degree functional language pathways [31, 34]. With auditory exposure and its learning processes via the dorsal pathways, infants may gradually construct the anterior-posterior language network in the first six months from birth, which may facilitate future language-related learning. This would eventually result in an enhanced role of SMG and STG for the processing of grammar-like structures. The results of Experiment 2 in 6-7-month-olds are consistent with this hypothesis.

### Activation in the posterior-language area for 6-7-month-old infants

Here we present the first evidence of specific brain regions engaging in auditory NAD processing in 6–7-month-olds, suggesting that the detection of NAD violations after learning may be subserved chiefly by anterior- and posterior-language areas (IFG, SMG, and STG) in addition to the comparable contralateral regions. This is consistent with findings in 9-month-olds processing syllabic NADs which engaged bilateral temporal regions [14]. Our results extend this finding firstly, by relating functional brain activation to the discrimination of NADs vs. NAD violations instead of NAD vs. no-NAD conditions [14] and secondly, by probing the brain basis of NAD learning at an even younger age. The activation pattern at 6-7 months is, to some extent, already the miniature of a mature adult brain activation in similar paradigms. Previous adult fMRI studies and reviews [48, 49] on artificial grammar learning have reported that frontoparietal areas, including left IFGoper, left inferior/superior parietal cortex, are involved in computing the syntactic regularity [28, 29, 50–52]. In addition, left posterior STG is often reported to be activated in these studies [51, 53, 54].

Activation of Brodmann’s area 44 (BA 44) has been reported to be associated with the processing of both rather simple as well as more complex linguistic structures [28, 29]. As we found activation in IFGoper (i.e., BA 44) and IFGtri (i.e., BA 45) in 6-7-month-olds in response to non-linguistic tone sequences, we suggest that inferior prefrontal regions play a core role in recognizing structural relations in the auditory environment after about half a year of extrauterine language exposure. The large STG response to Incorrect relative to Correct conditions may not only reflect the processing of physical auditory aspects of the stimuli but also computational aspects of processing. A previous ERP study showed that 4-month-olds can discriminate violation of NADs consisting of non-native syllables. These results were discussed in terms of the phonologically based associative learning mechanism mainly rooted in the temporal cortex although the respective experiments did not provide any functional anatomical information [15, 34].

Notably, we found activation in the bilateral SMG (one part of the inferior parietal lobe) in Experiment 2. The SMG has previously been associated with phonological working memory processes [55, 56] which also played a central role in a word segmentation task by 7 to 10-month-old infants [57]. During the Test phase in the present study, infants may have relied on auditory working memory processes in a similar way. Infants may generate memory-based expectations and their SMG responded upon coming across a violation, namely when the retrieval outcome contradicts those expectations [58]. It is possible that the activation in SMG, as a hub of the frontoparietal attention network, treats incorrect tone sequences presented in the Test phase as salient stimuli due to their unpredictable nature. This could result in infants involuntarily allocating their attention to these salient stimuli [59]. Such an interpretation is consistent with predictive coding theory, which assumes that internal forward models play an important role in many cognitive processes. Both STG and SMG have been suggested as playing a role during the processing of auditory sensorimotor predictions including in language production [60] and comprehension [61]. The activations found for the infants in the present study are consistent with the idea that predictions about upcoming tones in NAD sequences were formed and, in the case of violations, treated as prediction violations.

### NAD learning as an innate ability with ancient phylogenetic roots

Over and above the finding of specific brain regions involved in NAD learning and processing across development, our study demonstrates that this ability may be present in humans starting from birth onwards. It can be found for non-linguistic stimuli and it does not depend on a mature language network. Previous studies have also demonstrated this ability in both closely and distantly related species including apes, monkeys, rats, and birds [17, 62, 63]. While the basic ability to learn NADs may thus be present from early on, both ontogenetically and phylogenetically, the extensive use that human language makes of such structures may further shape both the cognitive processes and the neurophysiological substrate that is recruited during NAD processing. Such a view is congruent with human studies showing that the learning of simple NADs and even more complex sequential patterns varies as a function of the developmental status of learners [13, 64] and the way the patterns are learned [11]. The present findings may be taken to speculate that language development in the domain of sequential structure piggybacks on infants’ innate abilities to detect adjacent and even non-adjacent auditory patterns which may form an important basis for later syntactic development.

### Limitations

Deactivation was found in both experiments when comparing correct tone triplets with baseline (i.e. standard stimuli). The mere presence of a difference suggests that novel correct stimuli could be differentiated from familiar correct ones based on acoustic differences (i.e., pitch). As both experiments further yielded activation increases for novel incorrect stimuli compared to novel correct ones we must conclude that the underlying NAD rule led to a further differentiation of the two activation patterns. Note that the lack of a difference between novel incorrect vs. baseline stimuli in newborns is not informative about NAD detection alone, as this contrast includes both acoustic differences and a NAD violation, which may impact on the activity independently in different directions. Thus, we maintain the conclusion that both newborns and older infants extracted the NAD rule while acknowledging that the specific effects of the pitch variation and the rule violation yielded different effects across age.

Note, that we used nonlinguistic tones in our experiments in order to study processes that are especially important for language processing. The underlying aim was to minimize the effect of pre-natal language experience and processing load and to ensure comparability with animal research. While differences between tonal and speech stimuli have been reported during NAD learning in later infancy and early childhood studies [13], within the first half year of life rather report evidence for NAD learning in both domains [12, 15]. Furthermore, a review of developmental hemispheric lateralization in language acquisition [65] posits a model which assumes left-hemispheric learning biases for both speech and non-speech sounds. Thus, we assume comparability of basic involved mechanisms of linguistic and non-linguistic NAD learning in the first year of life while we acknowledge the need to further corroborate this in direct comparisons across modalities.

Another limitation of this study is that the infants’ states differed between Experiments 1 and 2. Measurements with neonates were performed during the active sleep state, but measurements with 6-7-month-old infants were performed during the awake state. Taga et al. [43] systematically examined the differences in brain activation and task-based FC with infants aged 2–10 months. They found that auditory stimuli produced a global activation for the sleep state but a focal activation for the awake state. In principle, activation foci were clearly different between our two experiments, namely the left prefrontal area for sleeping neonates and the left temporo-parietal area for the awake 6- 7-month-olds. These data imply sleeping state does not act as an obvious confound and hence should not impact the interpretation of our findings. However, the functional reorganization of brain regions supporting NAD learning across the first half year of life still needs to consider the state difference of infants in our two experiments.

To conclude, our study uses fNIRS to shed new light on the neurodevelopmental origins of a crucial building block of syntax - the capacity to process NADs. Specifically, we provide the first evidence that neonates are capable of extracting NADs from auditory sequences. Yet, neonates relied on different brain regions compared to 6-7- month-old infants on the same task. This suggests a rapid development of brain networks relevant to grammar-like processing within the first 6 months of life.

Specifically, 6-7-month-old infants already use the same brain regions (IFG and the posterior-language area) as adults to track NADs, while neonates recruited solely prefrontal regions. Neonates’ FCs during learning additionally revealed a learning-related brain network including posterior language-related areas that were found in 6-7- month-olds. These findings add to the growing evidence that the prefrontal cortex in concert with functionally connected posterior areas supports early cognitive development and indicates that the infant brain has already been equipped with an extraordinary learning ability as a precursor for later grammar acquisition.

## Materials and Methods

### Participants

For Experiment 1, 21 full-term healthy neonates (11 female, Mean age = 3.5 days, SD = 0.9, range: 1–5 days) contributed data to the final analyses. An additional two neonates were tested but were excluded due to fussiness/awake during the experiment. The number of neonates included in the functional connectivity (FC) analysis with valid FC data was 18, 17, 15 for the pre-task resting-state (Pre-Rest) phase, the learning (Learning) phase, and the post-task resting-state (Post-Rest) phase, respectively. All neonates had normal hearing which was assessed using auditory brainstem responses or other clinical tests. Mean birth weight for neonates was 2822 ± 358 g (range: 2078– 3527 g). For Experiment 2, 19 healthy 6-7-month-old full-term infants (6 female, Mean age = 201.2 days, SD = 9.8, range: 185–223 days) were included in the final analysis. Additional 22 infants were excluded from the analysis because of either rejection of wearing the fNIRS probe cap (n = 1), cessation of the experiment due to frequent fussiness (n = 12), or insufficient blocks caused by motion artifacts (n = 9). Written informed consent was obtained from parents before participation. Experiment 1 and 2 were approved by the ethics committee of Keio University Hospital (No. 20090189) and the ethics committee of Keio University, Faculty of Letters (Reference Number: 18009), respectively.

### Stimuli

The stimuli were identical to the materials used in a previous comparative study on NAD learning [17]. Each of artificial grammar sequences was composed of three elements from six computer-generated acoustic categories (A, B, C, D, X1, and X2) of frequency-modulated sine tones. Each category was characterized by its specific pitch contour (e.g., A: rising; B: slow wave; C: fast wave; D: falling; X1: big spike; X2: small spike; Fig 1A). The acoustic categories A, B, C, and D were composed of 10 pitch-shifted variants (SI Appendix, Table S1). The acoustic categories X1 and X2 consisted of 16 pitch-shifted variants (SI Appendix, Table S2). We created two sets of paired NAD grammars, i.e., Grammars 1 and 2. For Grammar 1, A elements were always followed by B elements (Grammar 1a), and C elements by D elements (Grammar 1b), whereas the middle intervening X elements varied freely. Conversely, the roles of B and D elements were reversed for Grammar 2, with D dependent on A (Grammar 2a) and B dependent on C (Grammar 2b). To control for the possibility that certain sound pairings might be relatively easier to learn, we created four stimulus lists, each of which consisted of the standard triplets, the correct triplets, and the incorrect triplets (SI Appendix, Table S3). The standard triplets and the correct triplets in one of four stimulus lists were used as one of two artificial grammars (e.g., Grammar 1) while the incorrect triplets as another one of two artificial grammars (e.g., Grammar 2). Further details are provided in SI Appendix, Stimuli Construction.

### Procedure

Experiment 1 was conducted in a testing room at the hospital and consisted of four phases, that is the Pre-Rest phase, the Learning phase, the Test phase, and the Post-Rest phase (Fig 1B and Fig 1C). In the Learning phase, neonates were presented with a total of 60 standard triplets (e.g., Grammar 1: 30 triplets for Grammar 1a and 30 triplets for Grammar 1b) drawn from one of the four stimulus lists (SI Appendix, Table S3) via two speakers positioned 45 cm from their heads while they were sleeping. While more than half of the sleeping period is in active sleeping state in neonates, they were in active sleeping states during the experiment as observed from visual inspection as well as the general time schedule of their daily care routine. Each triplet lasted 5 s and the time interval between two triplets was 1.5 s. Thus, 60 standard triplets resulted in the Learning phase lasting approximately 6 minutes. We used a block design in the Test phase such that the two types of target trials, Correct and Incorrect, were spaced by baseline trials varying duration (11.5 s, 18 s, and 24.5 s) to avoid synchronization between stimuli occurrences and spontaneous oscillations. The two target trial types were presented in a pseudorandom order, which was constrained to prevent more than two consecutive blocks of the same condition. The order of the trials was counterbalanced across neonates. A Correct trial was composed of two correct triplets, while an Incorrect trial included two incorrect triplets. Each triplet was 5 s and the time interval between two triplets was 1.5 s, meaning each target trial lasted 11.5 s in total.

For one given neonate, the stimuli for the Correct and Incorrect trials in the Test phase were selected from the same stimulus list as that presented in the Learning phase.

Assignment to four stimulus lists was counterbalanced across neonates as much as possible. The Test phase lasted for approximately 7.5 minutes since almost all neonates completed ten trials for each experimental condition. Besides, before the Learning phase and after the Test phase, two resting-state phases were incorporated to examine FC when neonates received no explicit tasks.

The procedure of Experiment 2 was slightly different from Experiment 1 in that the former consisted of only two phases, that is the Learning phase and the Test phase (Fig 1C). During the Learning and Test phases in Experiment 2, the same stimuli as Experiment 1 were presented to 6- 7-month-old infants in a sound-attenuated room. The infant sat on a mother’s lap and two loudspeakers were located approximately 1.5 m from the infant. An experimenter entertained the infant with silent toys, or a silent cartoon movie played on an iPad to avoid their fussiness. Additionally, the mother and the experimenter listened to music through headsets to prevent any potential influence on the infant’s behavior. After the Learning phase consisting of 60 standard triplets, two experimenters placed fNIRS probe pads onto the infant’s head. After the intensity level of all the channels had been checked, the Test phase immediately began. Here, we monitored the hemodynamic responses of infants exclusively during the Test phase to avoid the high attrition rate for infants at 6 months due to the long scan time. We ended the experiment after ten complete trials for each experimental condition, or when the infant became too fussy as judged by the experimenter.

### fNIRS measurements

The fNIRS measurements for Experiment 1 and 2 were conducted using two different multichannel NIRS instruments (ETG-4000, ETG-7000, Hitachi Medical Co., Japan) with the same sampling frequency of 10 Hz. Despite different probe pads, brain activities from frontal, temporal, and parietal regions were recorded according to the 10–20 system (SI Appendix, Fig S1). We utilized the virtual registration method [66] to estimate brain regions underlying each channel. Further details are provided in SI Appendix, fNIRS measurements.

### Data and statistical analysis

For Experiment 1: neonates, fNIRS data preprocessing was carried out on the MATLAB-based software Platform for Optical Topography Analysis Tools (POTATo, Hitachi Ltd., Research and Development) [67]. Based on the modified Beer-Lambert law [68], the optical density data were transformed into the product of oxygenated (HbO) or deoxygenated hemoglobin (HbR) concentration change and optical path length, which was defined as ΔHbO or ΔHbR (in mM⋅mm), respectively. Trials treated as contaminated by motion artifacts were excluded, according to the criteria for rapid signal change in the sums of ΔHbO and ΔHbR (>0.15 mM⋅mm changes within two consecutive data points). Trials with a saturated light intensity value (cf. more than 5 mM⋅mm) were also removed. For each neonate, data containing a minimum of two valid trials per condition and at least half of valid channels was included in the final data set. The time course of ΔHbO and ΔHbR then band-pass filtered using a third-order Butterworth filter between 0.02 and 0.5 Hz to eliminate slow drift and cardiac pulsation. Subsequently, the continuous data were segmented into blocks consisting of a 5-s baseline, 11.5-s experimental stimuli (Correct or Incorrect condition), and 11.5-s post-stimulus baseline. For these 28-s blocks, a baseline was linearly fitted between the first and last 5 s of each block. Lastly, we averaged all valid blocks for each channel per neonate. The average valid blocks across all neonates and channels were 6.9 (SD = 0.9) for Incorrect and 7.4 (SD = 1.5) Correct conditions, respectively. Group average waveforms from two experimental conditions (Correct and Incorrect conditions) were constructed over all neonates and channels to show the overall hemodynamic response shape so as to choose a time window of interest. Though a significant increase in ΔHbO or a significant decrease in ΔHbR could be regarded as an indicator of cortical activation, the most significant activation occurred in ΔHbO in previous studies [57, 69]. Thus, we restricted our statistical analysis to ΔHbO and conducted the statistical analysis using mean value of ΔHbO within the time window of 7.5s -12.5s from stimulus onset in this experiment.

For each channel, statistical comparisons (paired sample permutation t-tests) of average ΔHbO during the specified time windows were performed for the (1) Correct or Incorrect condition compared to the baseline condition and the (2) direct comparison of the Correct and Incorrect conditions. 50000 random permutations were implemented to estimate the distribution of the null hypothesis for an alpha level of 0.05.

In addition, we also performed the FC analyses. The pre-processing of continuous fNIRS data during the Pre-Rest, Learning, and Post-Rest phases was performed using the Homer2 package [70]. Detailed data preprocessing and artifact rejection are provided in SI Appendix, Preprocessing of FC data. If data included one “bad” channel due to motion artifacts or a saturated light within the continuous 90 s, it was excluded. Thus, the number of participants with valid FC data was 18, 17, 15 for the Pre-Rest, Learning, and Post-Rest phases, respectively. Finally, we extracted more than 90-s data (including more than 900 sample points) from continuous time courses of 46 channels for each neonate to perform further FC analyses.

In order to obtain significant FCs during the Pre-Rest, Learning, and Post-Rest phases, the first step of our analysis was to calculate the Pearson correlation coefficients (r) between the whole time course of HbO change from (46×45)/2 = 1035 pairs of channels for each neonate. We then converted r values to Z-values via Fisher’s r-to-z transformation to improve normality. One-sample t-tests were applied to find the significant FCs for each phase against a zero baseline. The cut-off levels of significance were set at 5 thresholds (p < 0.05, p < 5e-6, p < 5e-7, p < 5e-8, and p < 5e-9), and multiple comparisons among 1035 FCs were corrected using the Bonferroni method.

The second step of our analysis was to perform paired t-tests to determine FCs with significant changes relative to Pre-Rest by comparing FCs between Pre-Rest and Learning phases (Learning vs. Pre-Rest), as well as FCs between Pre-Rest and Post-Rest phases (Post-Rest vs. Pre-Rest). And when comparing FC differences for the contrasts of Learning vs. Pre-Rest and Post-Rest vs. Pre-Rest, the number of valid neonates was 15 and 14, respectively. The third step of our analysis was to examine whether there were significant associations between FCs with significant changes during/after NAD rule learning and the prefrontal activation engaged in NAD rule processing. We firstly chose 6 prefrontal channels (Ch30, Ch34, Ch35, Ch38, Ch39, and Ch43) that were significantly activated during the Test phase as seed channels, and then we calculated the Pearson correlation coefficients between the strength of FCs with significant changes (i.e., Learning minus Pre-Rest; Post-Rest minus Pre-Rest) and the degree of response in 6 prefrontal seed channels. For this analysis, only FCs which included any one of those 6 seed channels (i.e., seed-based FCs) were examined, because we aimed to find FCs involved in NAD learning (Fig 4A). The fourth step of our analysis was to examine whether these FCs formed a local brain network in response to NAD learning together with other FCs connecting any two non-seed channels (Ch 2, 9, 17, 19, and 22). Thus, we searched for FCs whose strength was significantly correlated with activation in 6 seed channels among all FCs with significant changes and simultaneously including non-seed channels (Ch2, 9, 17, 19, and 22). Given the local brain network underlying the NAD learning consisted of negative FCs, an additional analysis was performed to further explore the mechanism of the negative local brain network. Specifically, we analyzed the relationship between FC strength during the Pre-Rest phase (Pre-Rest minus 0) and that during the Learning phase (Learning minus 0), as well as the relationship between FC strength during the Pre-Rest phase (Pre-Rest minus 0) and FC strength changes from the Pre-Rest to Learning phases (Learning minus Pre-Rest). Likewise, among 19 FCs with significant changes from the Pre-Rest phase to Post-Rest phase, although we only found one FC which was negatively associated with the activity in the prefrontal channel, we also examined the relationship between FC strength during the Pre-Rest phase (Pre-Rest minus 0) and FC strength changes from the Pre-Rest to Post-Rest phases (Post-Rest minus Pre-Rest).

For the analysis of the task data collected during the Test phase in Experiment 2 (6-7- month-old infants), the data preprocessing and statistical analysis were identical to Experiment 1 (neonates), except that the mean value of ΔHbO within the time window of 10.5s -15.5s from stimulus onset was used for statistical comparisons according to overall hemodynamic response shape over all infants and channels. The average valid blocks across all infants and channels were 7.0 (SD = 2.1) for Incorrect and 7.1 (SD = 1.9) Correct conditions, respectively.

## Acknowledgments

We acknowledge all the staff in the Division of Neonatology of Keio University Hospital. We would like to thank all the infants and their parents who participated in this research. We also appreciate the assistance offered by Daisuke Tsuzuki, Sayaka Ishii and Hideka Osawa.

## Author Contributions

S.W., S.T., J.M., and Y.M. designed research; L.C., Y.H. M.H., T.A., N.S. and Y.M. performed research; L.C., E.H., and Y. M. analyzed data; and L.C., T.A., T.T., S.W., S.T., J.M., and Y.M. wrote the paper.

